# Dimensionally traceable 3D microstructures for multimodal microscope calibration

**DOI:** 10.64898/2026.05.07.722194

**Authors:** Junqing Jiang, Christopher W. Jones, Barry Reid, Dimitrios Tsikritsis, Ken Mingard, Poonam Ghai, Moona Kurttila, Michael Shaw

## Abstract

High-resolution microscopy techniques are used across research and industry to analyse biological systems, from biomolecules to subcellular organelles, multicellular models and tissues. As multimodal imaging workflows and quantitative analysis of bioimaging data become increasingly widespread, there is a growing need for materials and methods to calibrate imaging systems and evaluate the fidelity of generated image data. Here, we present three-dimensional microscopy phantoms fabricated using two-photon photolithography from transparent resins that exhibit both broadband visible autofluorescence and Raman scattering across the fingerprint and C-H stretching regions. Suitable for analysis using optical profilometry, the phantoms were dimensionally calibrated with SI traceability using a metrological confocal microscope. Immersible in air and common aqueous imaging media, the phantoms are compatible with a wide variety of optical microscopy techniques, including one and two-photon excited fluorescence and coherent Raman scattering microscopy. We employed a forked wedge design to validate image deconvolution results and a stacked lattice phantom to recover image distortion matrices under realistic biological imaging conditions. We demonstrate the impact of correcting chromatic offsets and axial scaling errors for a representative application: analysis of a cell seeded scaffold using confocal laser scanning fluorescence microscopy. These phantoms provide a versatile platform for calibration, quality control and validation of multimodal imaging pipelines and improved quantitative optical microscopy.

## INTRODUCTION

Light microscopy, particularly fluorescence microscopy, has long been the most widely-used form of high-resolution biological imaging. Live cell compatibility, high-spatiotemporal resolution, hardware accessibility and the availability of highly-specific (biomolecular) probes, give optical techniques unique advantages compared to other biological imaging modalities. New imaging techniques^1^, automated image acquisition workflows^2,3^, and widespread access to high-performance computing hardware and computational image analysis tools^4^ have created an unprecedented capacity for large scale capture and analysis of microscopy image data, which promises to increase the statistical power of bioimage-based analysis. The increased use of quantitative image analysis methods has created a corresponding need for improved calibration, standardization and quality control measures to ensure accuracy and reproducibility of results. National and international standards bodies, alongside community-based quality control initiatives^5^, are developing new documentary standards and protocols for characterizing key features of microscope performance^6^, as well as tools and guidelines for the analysis and reporting of microscopy image data^7^.

Physical calibration standards, reference materials and phantoms with well-defined structural and material properties are an important component of this drive to improve the quality and reliability of biomedical image data. Two-dimensional (2D) chrome on glass targets or etched substrates containing USAF line features and Siemens star designs are used to validate lateral spatial resolution in brightfield, coherent^8^ and phase contrast microscopy^9^, whilst grids and point arrays are used to measure geometric distortions. Such features are extruded vertically and combined with periodic structures and step height standards to quantify lateral resolution and calibrate 3D dimensional scales in surface topography measuring systems^10^. In fluorescence microscopy, fluorescent microspheres are widely used to measure instrument point spread functions and quantify spatial resolution^11,12^. Microspheres can be encapsulated within hydrogel matrices to create multimodal imaging phantoms for evaluating volumetric imaging performance under realistic biological imaging conditions^13^. DNA origami has been used to create ‘nanorulers’^14^ in which pairs of fluorophores are separated by a well-defined distance. However, sparse, randomly distributed features are unsuitable for measuring geometric distortions. Direct laser writing techniques have been used to create structured fluorescent calibration targets, comprising features (lines, points and uniform panels) within transparent polymer^15^ and glass substrates^16^. Whilst suitable for multichannel single photon excitation fluorescence microscopy, these targets are incompatible with many commonly used optical imaging techniques, including multiphoton and harmonic generation microscopy (which require high excitation irradiance) and Raman methods (which rely on chemical contrast between the processed and unprocessed substrate). Whilst all of these solutions allow assessment of imaging performance under idealized conditions, their dimensional characteristics and optical properties differ significantly from typical biological samples, limiting commutability to the real imaging context. Further (with some exceptions^15,17^) the targets are incompatible with optical dimensional metrology systems, which require well-defined surfaces, a large refractive index contrast, simple, extended geometrical features (such as flat facets and uniform, repeating periodic structures), and minimal shadowing, for accurate, traceable calibration.

Two photon photolithography (TPP) can be used to fabricate high resolution 3D microstructures within transparent substrates^18^ and has recently been applied to create phantoms for quantitative phase imaging^19^. This article describes our work to apply TPP to create commutable, multimodal optical microscopy calibration phantoms from commercially available transparent photoresins. We present three scalable designs and investigate fabrication results using scanning electron microscopy (SEM). The spectroscopic properties of different cured resins are analysed to establish compatibility with fluorescence and Raman microscopy. We then image the same pair of phantoms using confocal laser scanning fluorescence microscopy (CLSFM), light sheet fluorescence microscopy (LSFM), two-photon excited fluorescence microscopy (TPEFM) and stimulated Raman scattering microscopy (SRSM) platforms. We traceably calibrate the topography of a forked wedge and multilayered grid phantoms using a metrological optical profilometry system and use these phantoms to validate image deconvolution results in LSFM and derive a set of 3D affine transformations for correcting chromatic offsets and axial distortions in a CLSFM system. After verifying correction results, these transforms are used to correct images of a 3D cell-seeded scaffold. Differences between the raw and uncorrected images serve to illustrate the impact of dimensional image calibration on the interpretation of image results. Finally, we discuss the compatibility of the phantoms with additional imaging modalities and consider further development and exploitation of the core concept to support other quantitative and multimodal bioimaging workflows.

## RESULTS AND DISCUSSION

### Design and verification of imaging phantoms

We took inspiration for the design of 3D phantoms from existing 2D calibration targets including rectilinear grids and line pairs, and established methods for traceable measurement of areal surface topography^20^. Grid structures are well-suited for measurement of geometric field distortions, whilst line pairs with different spacings enable assessment of lateral spatial resolution. After initial optimisation, three phantom designs meeting design criteria were fabricated on glass coverslips using two-photon photolithography. The ‘stacked lattice’ phantom (Fig. 1(a)) comprised three rectilinear lattices stacked on top of each other. Intersections, edges and corners in each layer were designed to provide fiducial points for calibration and image registration, whilst the 35° rotation between consecutive layers allows direct visualisation of features in deeper layers without occlusion by preceding layers, essential for compatibility with the optical profilometer used to calibrate the phantoms. Comprising line pairs spaced by between 0.8 µm and 12.0 µm, the ‘forked wedges’ (Fig. 1(b)) were designed for assessing lateral spatial resolution as a function of axial position. The wedge slope angle provides a feature for assessing optical sectioning capability and axial scaling. Finally, the ‘cylinder array’ (Fig. 1(c)) comprised 36 tubes of varying height and wall thickness to evaluate both lateral spatial resolution (line spread function) and axial scaling. All designs were formed from hollow elements with simple geometries and minimal overhanging features, making them suitable for fabrication using 2-photon photolithography.

**Figure 1.**
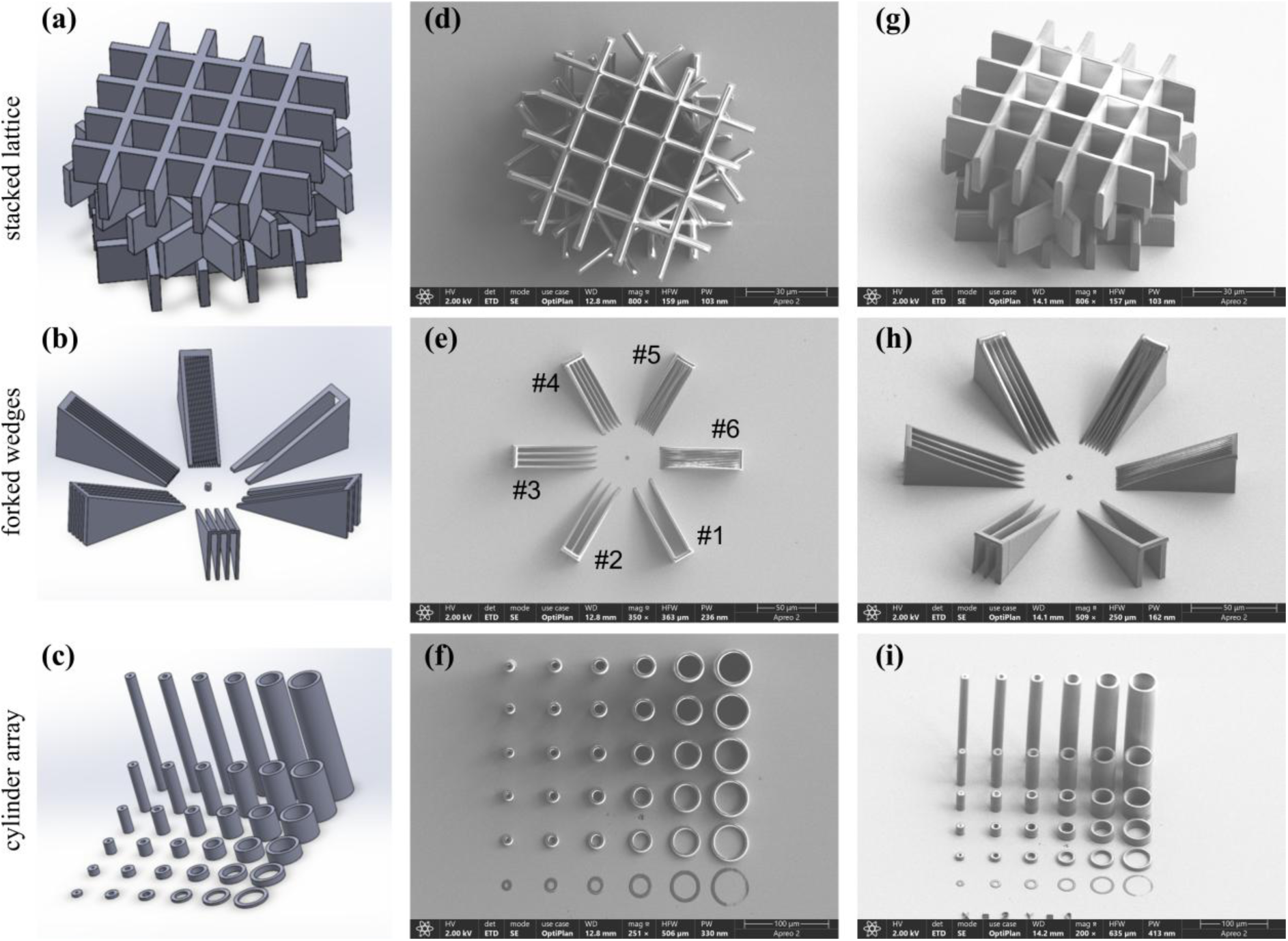
Design and validation of 3D calibration phantoms. (a-c) CAD models for stacked lattice, forked wedges and cylinder array calibration targets. En face (d-e) and oblique (g-i) SEM micrographs of phantoms fabricated using TPP.

A series of phantoms were fabricated at different scales – with a lateral footprint from 100 μm to 500 μm - from two commercially available resins (IP-L and IP-S) and imaged using scanning electron microscopy (SEM) for qualitative structural verification. Figure 1(d-i) shows representative micrographs of each design made from IP-S, similar results were obtained for phantoms fabricated at different sizes and for both resins (see supplementary Fig. S1). Images of the stacked lattice (Fig. 1(d,g)) confirm that all three layers were fabricated successfully with minimal layer-to-layer merging. Fabrication was also successful for the low pitch forked wedges (Fig. 1(e,h)). However, wedges #5 and #6 were not mechanically stable with internal panels collapsing and merging - likely during post fabrication development. Images of the cylinder array (Fig. 1(f,i)) show fabrication errors for the shortest cylinders (2.4 µm). The oblique view (Fig. 1(i)) also indicates that wall thickness varies along axial direction. The limited contact area with the substrate meant that taller cylinders had a tendency to collapse. As a result, only the stacked lattice and forked wedge phantom designs were taken forward for further testing and analysis.

### Spectroscopic characterization

To investigate the compatibility of the 3D imaging phantoms with fluorescence and Raman microscopy systems, we performed spectroscopic analysis of both photoresins. Figure 2(a) shows fluorescence emission spectra of cured IP-S and IP-L under excitation at four wavelengths commonly used in fluorescence microscopy: 365 nm (used to excite DNA binding dyes including DAPI and Hoechst), 488 nm (used to excite green fluorescent proteins and fluorescein-based dyes) and 561 nm (used to excite red fluorescent proteins and rhodamine derivatives). Both resins exhibited measurable autofluorescence at all excitation wavelengths. However, the fluorescent intensity was significantly greater under short wavelength (365 nm and 488 nm) excitation. Significant differences in the shape of the spectrum and the position of the emission maxima were observed for the two resins, although the relatively broadband emission indicates that both are compatible with common fluorescence filter sets. Fluorescent emission was also observed under excitation at 638 nm (a wavelength used to excite far-red fluorescent dyes including Cyanine5 and Texas Red), however the low fluorescence quantum yield combined with the small Stokes shift prevented reliable measurement of the emission spectrum.

**Figure 2.**
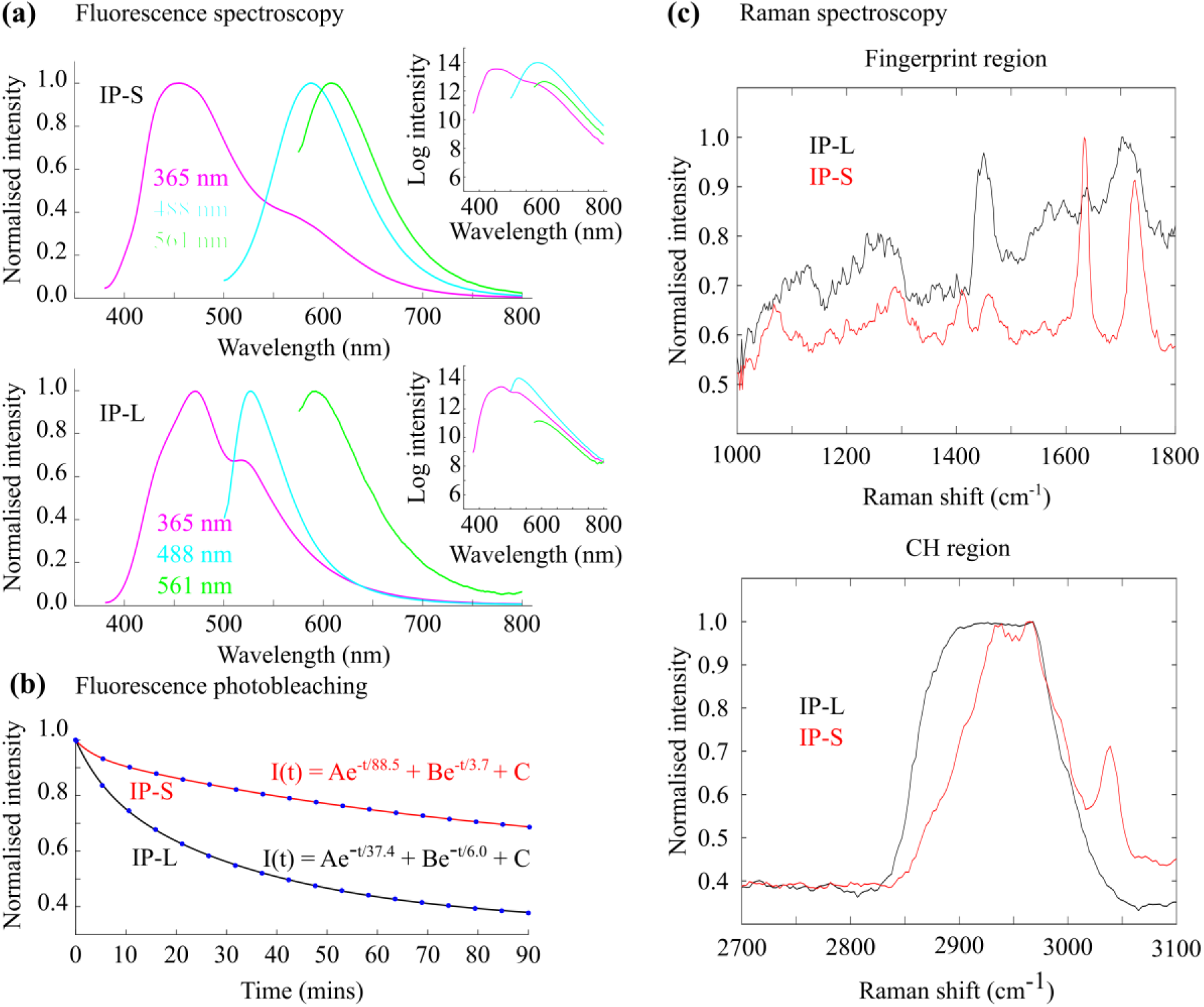
Fluorescence and Raman spectroscopic characterization of cured IP-S and IP-L resins. (a) Fluorescence emission spectra of IP-S and IP-L resins at three fluorescent excitation wavelengths commonly used in fluorescence microscopy. (b) Intensity decay of emitted fluorescence intensity from cured resin under irradiance at 54 mWcm^−2^ at 488 nm. (c) Raman spectra of the two photoresins in the fingerprint (left) and CH (right) regions.

To assess photostability we measured the changes in emitted fluorescence when both resins were exposed to 488 nm illumination at 54 mWcm^−2^. IP-S proved more stable than IP-L, with fluorescence intensity decreasing to 74% of its initial value following an hour of continuous exposure, whereas for IP-L the emitted intensity was reduced to 44% after the same period.

Accurate fitting of the decay curves required a two-component exponential (Fig. 2(b)) comprising one rapidly decaying term with a short half-life (2.6 mins and 4.2 mins for IP-S and IP-L respectively) and a slowly varying term with a longer half-life (61 mins and 26 mins for IP-S and IP-L respectively) - suggesting two discrete fluorescent transitions.

Both resins exhibited significant Raman scattering (Fig. 2(c)) across the fingerprint and C-H stretching regions, commonly analysed in spontaneous and coherent Raman scattering microscopy techniques. Differences between the spectra may reflect chemical differences or different degrees of polymerisation. IP-S has prominent peaks at ∼1636 cm^−1^ (indicating carbon-carbon double bonds), 1725 cm^−1^ (associated with ester groups) and 3041 cm^−1^ (associated with CH stretches). Both spectra contain broad peaks from ∼ 2900 cm^−1^ to 3000 cm^−1^, typical of CH, CH_2_ and CH_3_ stretching modes of aliphatic chains.

### Optical profilometry of imaging phantoms

The surface topography of stacked lattice and forked wedge phantoms was measured in air using an optical profilometer. Figure 3(a) shows a reflected light confocal intensity image of a 200 µm stacked lattice phantom fabricated from IP-L; the corresponding areal topography map and pixel height histogram with respect to the surface of the coverslip are shown in Figure 3(b). Analysis of histogram centroids indicates that the top surfaces of the three lattice layers are 33.9 µm ± 0.9 µm, 66.9 µm ± 0.7 µm, and 100.3 ± 0.9 µm above the substrate – consistent with the design values of 33.72 µm, 67.44 µm, and 101.16 µm. Local flatness variations across the surface of each layer contribute to the spread of measured height values (the full width at half maximum of the height histograms for each layer are 1.0 µm, 0.6 µm and 0.5 µm) and a slight asymmetry in the histograms. Although comparable in magnitude to the axial resolution achievable using high NA oil immersion objective lenses, these flatness deviations are significantly less than the nominal axial resolution of the air and water immersion objective lenses used to image the phantoms in this work. The lateral dimensions of the top lattice layer were characterised by the distances between the lattice crossing points along the cartesian axes nominally aligned to the lattice bars. By 2D cross-correlation of a 15 µm x 15 µm intersection template (Fig. 3(c)) over the height map, the mean spacing between 16 lattice intersection points (Fig. 3(d)) was measured as 33.09 µm ± 0.17 µm in local X and 33.39 µm ± 0.13 µm in local Y, compared with a design grid pitch of 33.72 µm (the sum of the 6.74 µm bar width and 26.98 µm clear aperture).

**Figure 3.**
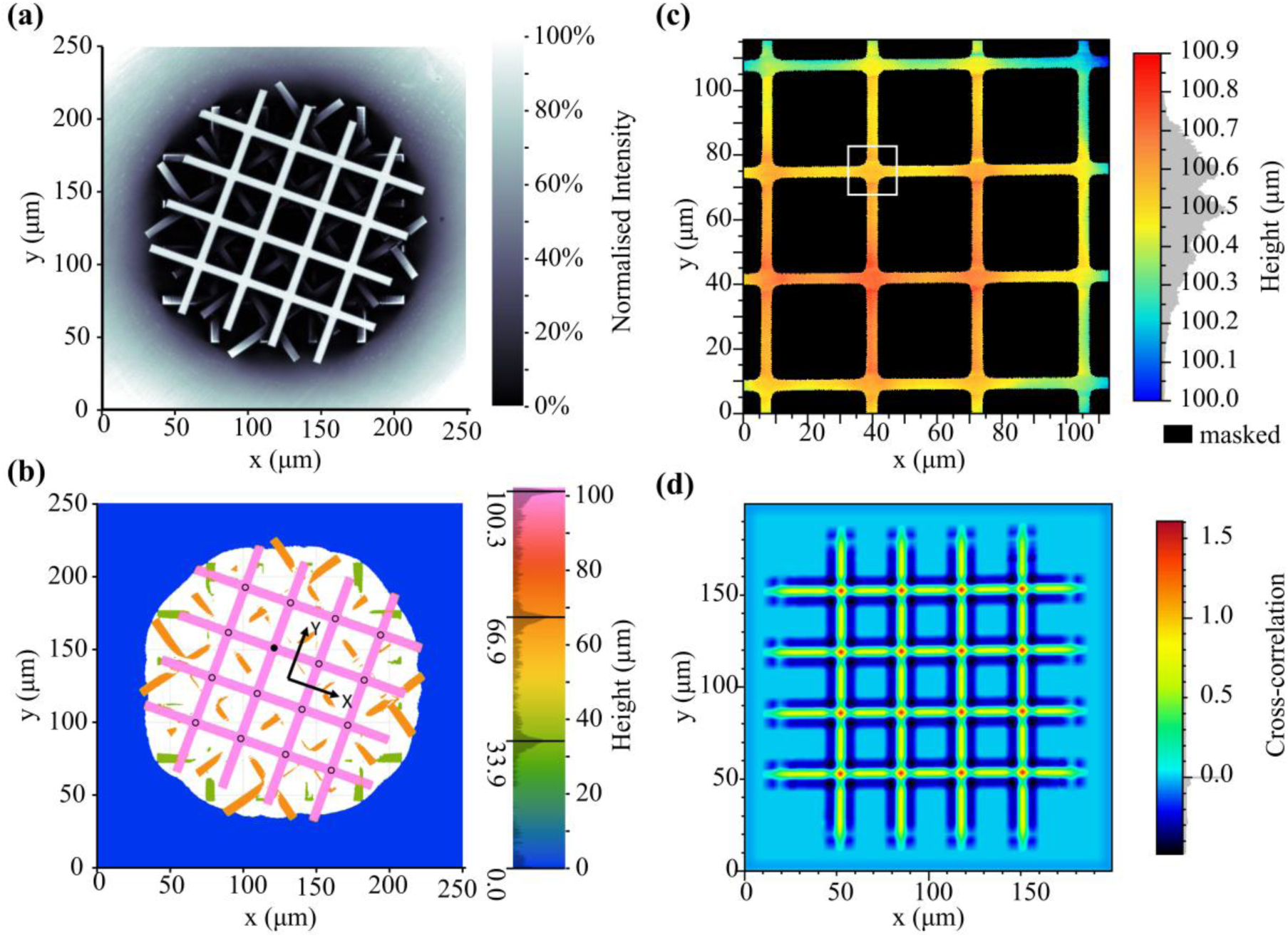
Dimensional calibration of a stacked lattice phantom. (a) Reflected light confocal microscopy intensity image of a stacked lattice phantom. (b) Masked surface topography image for the same phantom highlighting the distance from the substrate to the top of each lattice layer. (c) Height map for the top lattice layer with an intersection template highlighted by the white square. (d) Two dimensional cross-correlation of the intersection template over the height image. The 16 local maxima at grid bar intersections define a set of fiducials for characterizing the lateral grid spacing.

The forked wedge phantoms contain fixed-angle features, enabling a test of optical sectioning and a check of the lateral and vertical axes. Apparent deviation from the calibrated wedge angle suggests a scaling of XY and/or Z axes, whilst deviations from constant slope suggest non-linearities in at least one axis. A least squares linear fit to line profiles drawn along the prongs of the leftmost wedge in the 400 µm-sized forked wedge phantom (Fig. 4) gave an estimated wedge angle of 24.4° ± 0.6°, consistent with the design of 25°, with an associated linearity deviation of 0.5 µm.

**Figure 4.**
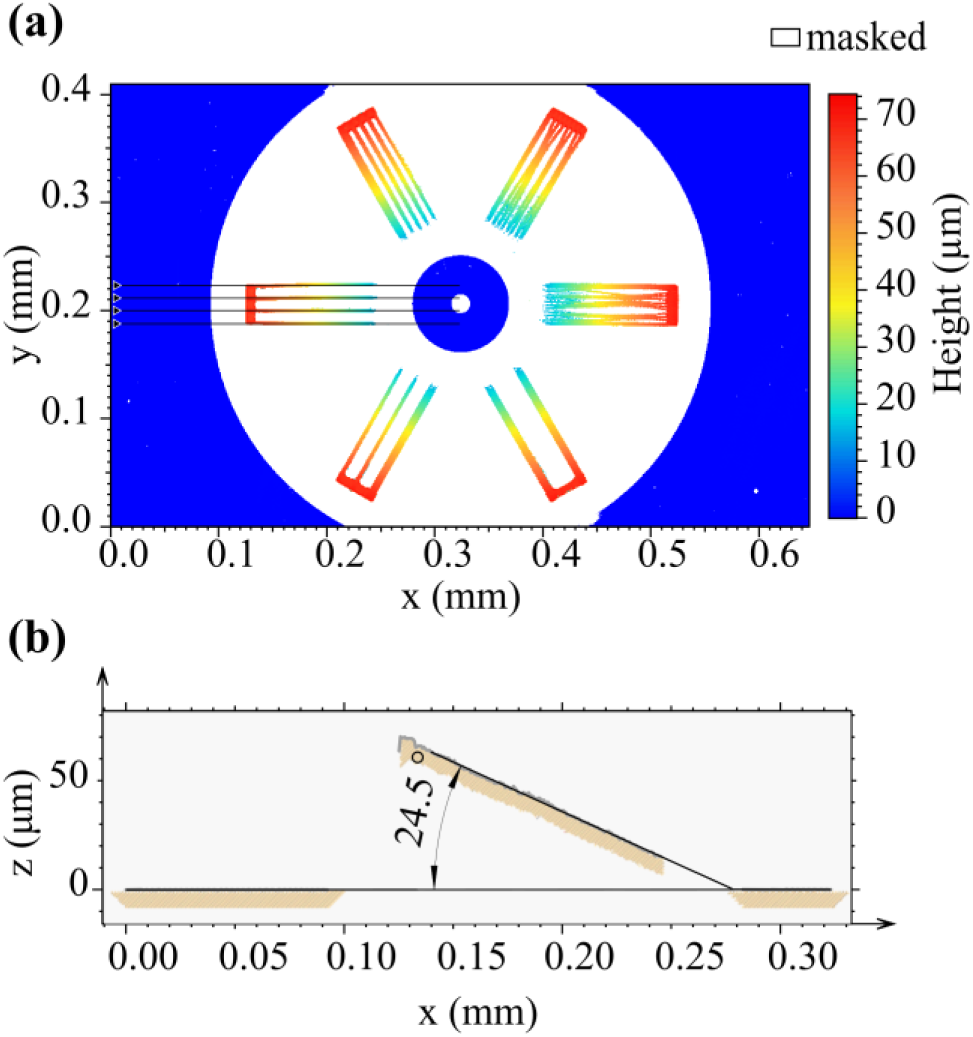
Dimensional calibration of a forked wedge phantom. (a) Surface topography of the phantom (fabricated from IP-S) measured using optical profilometry. (b) Wedge angle for the leftmost fork calculated from the line profiles shown in (a).

To assess structural stability, the topography of a prototype stacked lattice was measured twice, with a period of three months between measurements during which the sample was stored in a microscope slide holder in a temperature-controlled laboratory. Comparison of the two sets of measurement data found no significant differences in axial and lateral feature spacings quantified as described above (see supplementary Fig. S2).

### Fluorescence and stimulated Raman scattering microscopy of phantoms

Stacked lattice and forked wedge phantoms were imaged using three commercial microscope platforms: a confocal laser scanning fluorescence microscope (CLSFM); a stimulated Raman scattering microscope (SRSM), also capable of two photon excited fluorescence microscopy (TPEFM); and an Airy beam light sheet fluorescence microscope (LSFM). For the inverted CLSFM, SRSM and TPEFM systems, phantoms were imaged through the coverslip, whilst LSFM images were captured from above using a matched pair of water immersion objective lenses. In all cases, images were captured using instrumental configurations (irradiance, beam scanning and detector settings) typically used for biological samples. Qualitative analysis of CLSFM, SRSM and TPEFM results (Fig. 5(a-c)) indicates images are free from obvious artefacts, such as geometric distortions, or striping/patterning due to intensity fluctuations or beam scanning errors.

**Figure 5.**
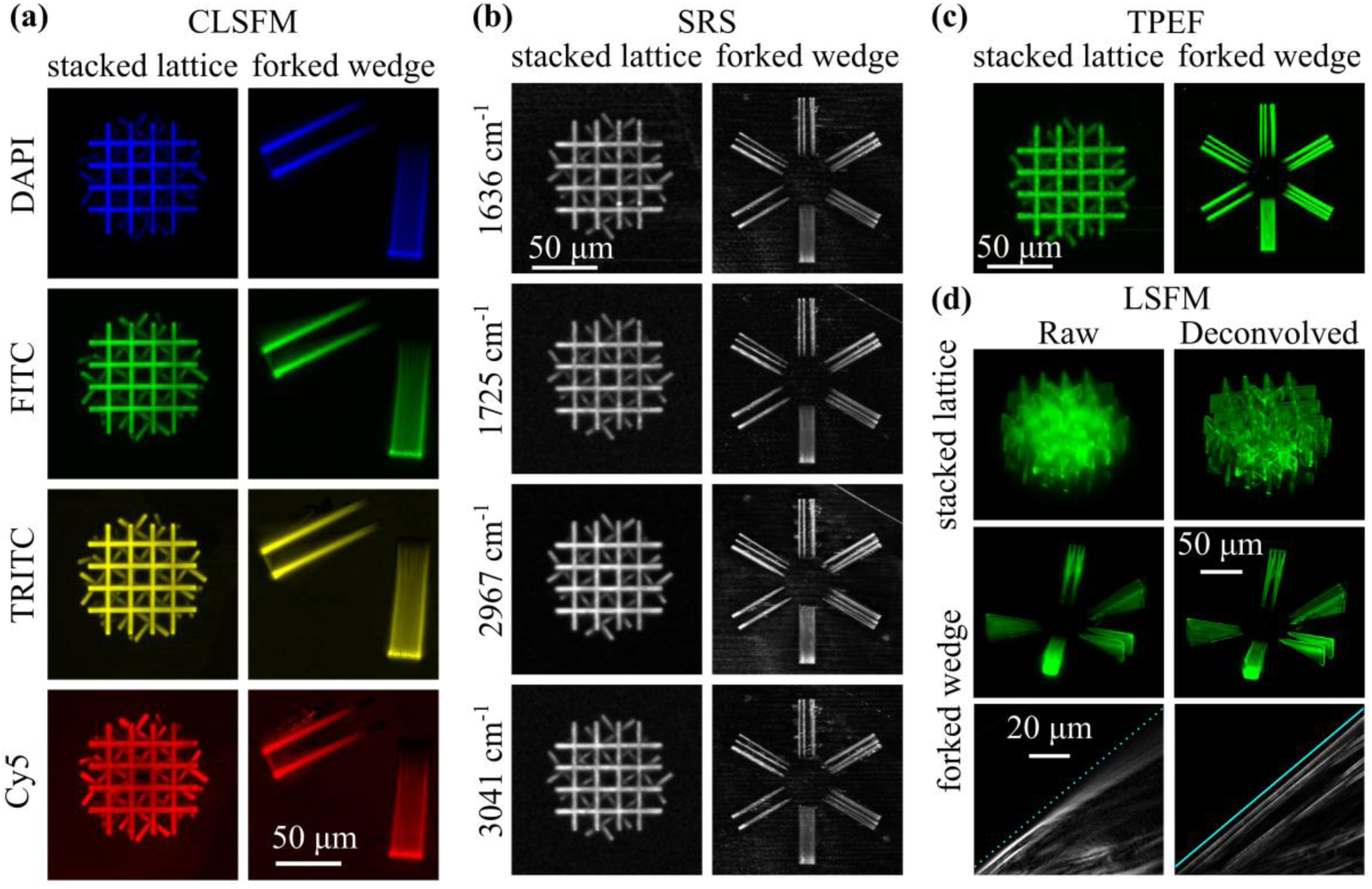
Multichannel images of the same stacked lattice and forked wedge phantoms captured using four common biological microscopy techniques. (a) CLSFM images in four fluorescence channels. (b) SRSM images at wavenumbers close to the vibrational modes of four common molecular bonds. (c) TPEFM images of the same phantoms. CLSFM, SRSM and TPEFM images are maximum intensity projections along the axial direction of a focal series. (d) LSFM images before (left) and after (right) Richardson-Lucy deconvolution. The straight lines (cyan) superimposed onto the lowest pair of images show a (vertically offset) linear least squares fit to the top surface of the wedge in the deconvolved image.

Well-defined geometrical features are particularly useful for verifying Airy beam LSFM results. Airy beams follow a curved trajectory, with a transverse intensity profile containing a series of side lobes in addition to the main intensity maximum^21^. As a result, raw Airy beam LSFM images are typically processed using 3D deconvolution techniques to correct image warping and shadowing artefacts. Figure 5(d) shows Airy beam LSFM images of stacked lattice and forked wedge phantoms before (left) and after (right) deconvolution using the Richardson-Lucy algorithm^22,23^ with an empirical point spread function (PSF). Deconvolution significantly increases visual image contrast for both phantoms. To validate the correction of the warped image volume due to the curvature of the Airy beam, we analysed the surface of one of the wedges calibrated previously using optical profilometry. The cyan lines superimposed on the grayscale images in Fig. 5(d) show a least squares linear fit to the surface of the wedge in the deconvolved image (lines have been offset vertically so the intensity features are visible). Whereas in the raw image the wedge surface appears curved, after deconvolution the surface is accurately fit by a first order polynomial (R-squared = 0.9999).

To further demonstrate further practical application of the phantoms, CLSFM images of a 100 µm stacked lattice (fabricated from IP-L) immersed in water were used to measure chromatic offsets between DAPI, FITC and Cy5 channel images captured using a 20x/0.75 objective lens. Chromatic correction matrices were derived from the image data based on lateral displacement of 48 features (Fig. 6(a)) between the different fluorescence channels. A set of 45 x 45 pixel template patches centred on features in the DAPI channel were cross-correlated with the FITC and Cy5 images, with the maximum correlation defining the feature location in each channel. The resulting offsets (Fig. 6(b)) were used to derive a pair of 3D affine transformation matrices to map FITC and Cy5 data to the DAPI channel image space. Chromatic corrections were checked using images of multicolour fluorescent microspheres (Tetraspeck, 0.5 µm, Thermofisher) immobilized in a 1% agarose gel. In uncorrected data the maximum/mean lateral offsets between bead centroids in FITC-DAPI and Cy5-DAPI channels were 985 nm/876 nm and 1179 nm/1016 nm respectively (Fig. 6(c)). After correction, these offsets were reduced to 872 nm/622 nm and 584 nm/481 nm.

**Figure 6.**
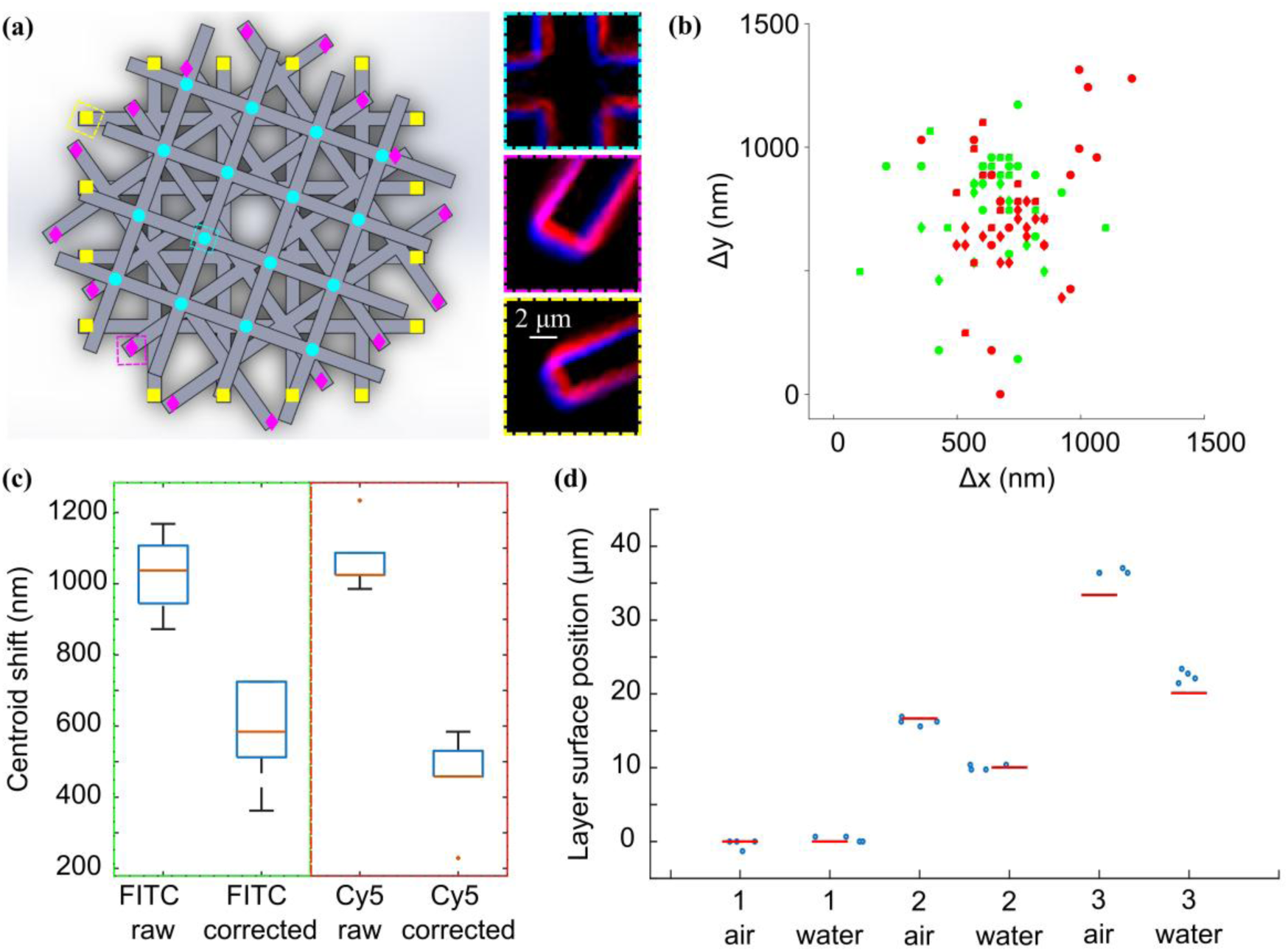
Measurement and correction of chromatic offsets and focus shifts in CLSFM using a stacked lattice phantom. (a) CAD rendering of the phantom with ROI centres used to determine lateral chromatic offsets highlighted. Zoomed views to the right of the main image panel show ROIs from each layer imaged in DAPI (blue) and Cy5 (red) channels. (b) Measured lateral offset in FITC (green), and Cy5 (red) channels relative to DAPI channel for the 48 features used to derive the 3D chromatic distortion matrix. (c) Box whisker plots showing centroid offsets between DAPI and FITC (green box) and DAPI and Cy5 (red box) for multicolour PSL particles before (left) and after (right) correction. (d) Measured layer spacing (circles) derived from axial intensity profiles in images captured using a dry objective lens for the phantom immersed in air (left) and water (right). Red lines show layer positions based on topographical calibration in air and a geometric model of focal shift due to air-water refractive index mismatch.

The same phantom was used to measure axial distortion of the image volume caused by a mismatch between the refractive index of the objective immersion medium and the sample. This phenomenon has been extensively studied in optical microscopy and several analytical models have been proposed for correction^24^. Containing well-defined planes which span an axial range exceeding that used in most high-resolution fluorescence imaging applications, the stacked lattice can be used to measure focal shifts under different experimental conditions. Such an empirical assessment is particularly useful if the sample refractive index and optical properties of the microscope are not accurately known. As a demonstration we imaged the phantom in air and water using the same (20x/0.75) dry objective lens. The 100 µm stacked lattice has a nominal layer spacing of 16.7 µm with calibrated spacings of 16.67 µm and 16.73 µm between the first and second and second and third layers respectively. Based on the separation of maxima in axial intensity profiles, when immersed in air the measured distances from the first to the second and second to third layers were 16.6 µm and 20.3 µm - close to these nominal values (Fig. 6(d)). However, when immersed in water the focus shift due to the refractive index difference resulted in axial compression of the imaging volume, with the separation between the first and second and second and third surfaces appearing to be 9.8 µm and 12.3 µm. These shifts are in excellent agreement with the focus shift predicted using the geometrical model described by Visser et al.^25^, 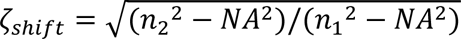

To quantify the magnitude of lateral offsets between SRSM and TPEFM image data, the scale invariant feature transform (SIFT) algorithm^26^ was used to identify common features in (4x) upsampled maximum intensity (z-projections) of SRS and TPEF images of the 100 µm stacked lattice phantom. The maximum offset between corresponding keypoints in TPEF and SRS images (at 1636 cm^−1^, 1725 cm^−1^, 2967 cm^−1^, and 3041 cm^−1^) varied between 1.9 µm and 2.6 µm. These values are significantly larger than both the raw pixel size and nominal spatial resolution of the microscope and, if uncorrected, would contribute significantly to errors and uncertainties in co-localisation measurements.

### Dimensional correction of biological images

To demonstrate the combined effect of focus shifts and chromatic offsets on biological image data, transformation matrices were applied to correct images of GFP expressing HeLa cells cultured in a 3D collagen scaffold captured using a 20x/0.75 air immersion objective lens. Affine transformation matrices derived from images of a 100 µm stacked lattice phantom were modified to correct for the axial scaling due to the refractive index mismatch between the objective immersion media (n = 1) and the specimen (n ∼ 1.33). As illustrated in Figure 7(a), at the cellular scale even relatively small geometrical errors can have a significant effect on the interpretation of image data. The zoomed regions of interest on the right of the main figure panel show an individual cell nucleus (blue) and several small vesicles (green). In the uncorrected image (top) a pair of bright vesicles appear in close proximity to the periphery of the nucleus. Based on the intensity distribution, the vesicles could be classified as either in contact with the nuclear membrane or inside the nucleus - although the low image contrast and spatial-resolution make it difficult to make a definitive assessment. However, the corrected image (below) reveals that the vesicles are clearly separate from the nucleus. This is further highlighted by the intensity line profiles below the figures. In the uncorrected image, the intensity maximum associated with the leftmost vesicle coincides with significant signal in the DAPI channel, suggesting nuclear co-localisation. Following geometric correction of the image volume, the same vesicle is displaced away from the nucleus locating it over a background region in the DAPI channel. Figure 7(b) shows the GFP channel for a different region of interest in the same sample before (magenta) and after (cyan) correction. The lateral displacement of both GFP positive vesicles and cell bodies, following correction is clearly visible.

**Figure 7.**
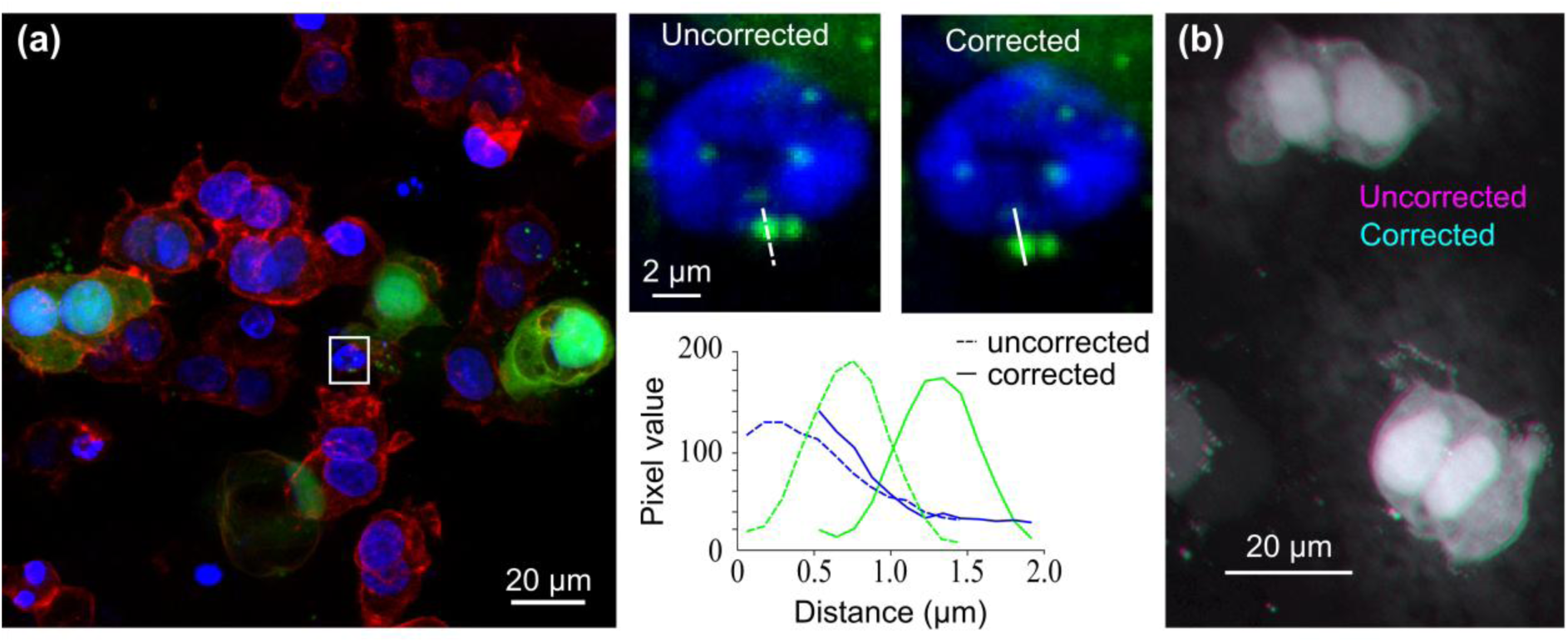
Correction of multichannel confocal fluorescence microscopy image data using phantom-derived chromatic distortion matrices. (a) Maximum intensity (z) projection of collagen matrix seeded with GFP expressing HeLa cells (green) stained for DNA (blue) and F-actin (red). Zoomed views highlight changes to the location of GFP-stained vesicles relative to the nucleus of one of the cells following correction of geometric image distortions. The plot underneath shows the signal along the white lines in the zoomed views. (b) GFP channel before (magenta) and after (cyan) correction for a different region within the same sample.

## CONCLUSIONS

We have demonstrated that 3D microstructures fabricated using TPP can serve as phantoms for testing and dimensional calibration of a range of light microscope systems. Exhibiting strong autofluorescence and Raman scattering across spectral ranges commonly used in biological imaging studies, the phantoms are compatible with a range of fluorescence and coherent Raman microscopy techniques including CLSFM, LSFM, TPEFM and SRSM, making them particularly well-suited for use in multi-technology imaging facilities and for testing data integration and fusion methods in multimodal and correlative imaging workflows. Structurally stable and immersible in a variety of immersion media, the phantoms can be used to test imaging performance under realistic biological imaging conditions, validate image reconstruction and processing workflows and correct chromatic offsets and axial distortions arising from refractive index mismatches between the objective and specimen immersion media. As a result, we anticipate a potential application for 2PP phantoms in correcting image data from cleared tissues^27^ for which the refractive index of the clearing solution is often uncertain.

In addition to the fluorescence and Raman microscopy techniques described in this article, there are further bioimaging modalities for which the phantoms may be compatible. Secondary ion mass spectroscopy (SIMS)^28^ can achieve similar spatial resolution to optical microscopy and has been used previous in correlative imaging studies with fluorescence and Raman methods^29^. In preliminary testing, the high energy ion source in a Time-of-Flight SIMS system caused sample heating and subsequent deformation. However, further optimization of acquisition settings and testing with other hardware may reveal a use case for the phantoms in SIMS calibration and quality control. Other options to increase the utility of the phantoms for chemical imaging include post fabrication surface functionalization or the addition of small quantities of dopants to add specific chemical groups. Alternatively, cured phantoms could be immersed in solutions containing chemical groups of interest to create a negative calibration sample.

IP-S and IP-L proved suitable substrates for fabricating stable phantoms compatible with a range of imaging techniques, however after curing both resins have a significantly higher refractive index (∼1.51) than common biological samples – which typically have a refractive index in the range 1.34 to 1.42^30^. As a result, despite the transparency of the resins, it was necessary to rely on directly accessible (unobscured) features to derive correction matrices. Attempts at fabrication with other low refractive index polymers^31^ proved unsuccessful. However, further optimization, including the addition of a two-photon polymerization initiator could enable use of a wider range of resins with tailored optical or chemical properties. Further opportunities for development include alternative methods for extracting correction matrices. In particular, the method for estimating layer separation in the twisted lattice phantoms differed between optical profilometry and CLSFM data – potentially introducing systematic errors into the axial distortion correction.

Although fabrication requires two-photon photolithography hardware, which is typically only available at large industrial and research institutes, per unit costs for each phantom are negligible. Accounting for material wastage, several phantoms could be produced for less than £1. Their low cost combined with multiplatform compatibility and stability make the phantoms an accessible tool to support efforts to improve data quality in biological microscopy.

## MATERIALS AND METHODS

### Two-photon photolithography

Three-dimensional microstructures were fabricated by direct laser writing (DLW) using a Nanoscribe Photonic Professional GT+ system (Nanoscribe). This technique employs a femtosecond-pulsed laser (780 nm) focused within a photosensitive resin through a microscope objective lens. The femtosecond pulses generate sufficient energy to polymerise the photoresist at a focal spot due to the phenomenon of two photon polymerisation (2PP), achieving greater resolution and voxel control when compared to standard curing process using shorter wavelength light sources^32^. Laser beam steering in the lateral (xy) plane was achieved using a galvanometric scanner, while vertical (z-axis) positioning was controlled with a high-precision piezoelectric stage. To optimise print quality, a parameter sweep was conducted, in which laser power, slicing (in-plane) distance, hatching (out-of-plane) distance. Optimal settings were determined to be a laser power of 35 mW (70% of the calibrated 50 mW output), a slicing distance of 0.2 µm, and a hatching distance of 0.1 µm. Printing was performed in Dip-in Laser Lithography (DiLL) mode, with the objective lens immersed directly into the resin to accurately locate the interface between the resin and the substrate surface. Post-fabrication, residual, unpolymerised resin was removed by immersing the samples in propylene glycol monomethyl ether acetate (PGMEA, ReagentPlus grade, ≥99.5%; Merck Sigma-Aldrich) for 20 minutes, followed by a 5-minute rinse in 2-propanol (≥99.5%; Merck Sigma-Aldrich). Samples were then dried under a stream of nitrogen gas (zero grade; BOC).

Two proprietary negative-tone photoresists were evaluated: IP-S (methacrylate-based) and IP-L (acrylate-based). While IP-S is recommended by Nanoscribe for medium-resolution structures fabrication used a 25x objective, IP-L was also tested due to its potential for superior resolution and smoother surface finish. Substrates consisted of indium tin oxide (ITO) coated (#1.5) borosilicate glass coverslips. Prior to printing, coverslips were cleaned sequentially with acetone and 2-propanol, followed by oxygen plasma treatment (Henniker Plasma HPT-100, 100 W, 1 min) to enhance surface wettability and adhesion. A droplet of photoresist was deposited onto the treated substrate, which was then mounted onto the sample holder and inserted into the system.

To simplify sample mounting and handling for subsequent imaging, the coverslip was glued to the top of an aluminum plate cut to the approximate size of a standard microscope slide. A (30 mm x 14 mm) rectangular hole in the centre of the plate allowed optical access to the phantom from above and below. To image phantoms immersed in water using CLSFM, a water droplet was added directly to the phantom. For SRSM and TPEF imaging using an oil immersion condenser, a 200 µm thick silicon spacer ring was placed around the phantom. After the ring was filled with water a second #1.5 coverslip was added on top.

### Fluorescence and Raman spectroscopy

Fluorescence spectra of curved resins were measured using an FLS1000 Photoluminescence Spectrometer (Edinburgh Instruments). The sample was deposited on a strip of aluminium foil which was positioned in the sample holder at a 45° angle from both the excitation and emission arm. After positioning, the strip was secured in place and emission spectra were collected at room temperature with a 1 nm step size and 0.1 s/step dwell time. Emission spectra were collected at four different excitation wavelengths: 365 nm, 488 nm, 561 nm, and 638 nm. The spectrometer slit width was optimized for each excitation wavelength for best signal-to-noise ratio, and the measurement parameters for each are optimized in Table 1.

**Table 1.**
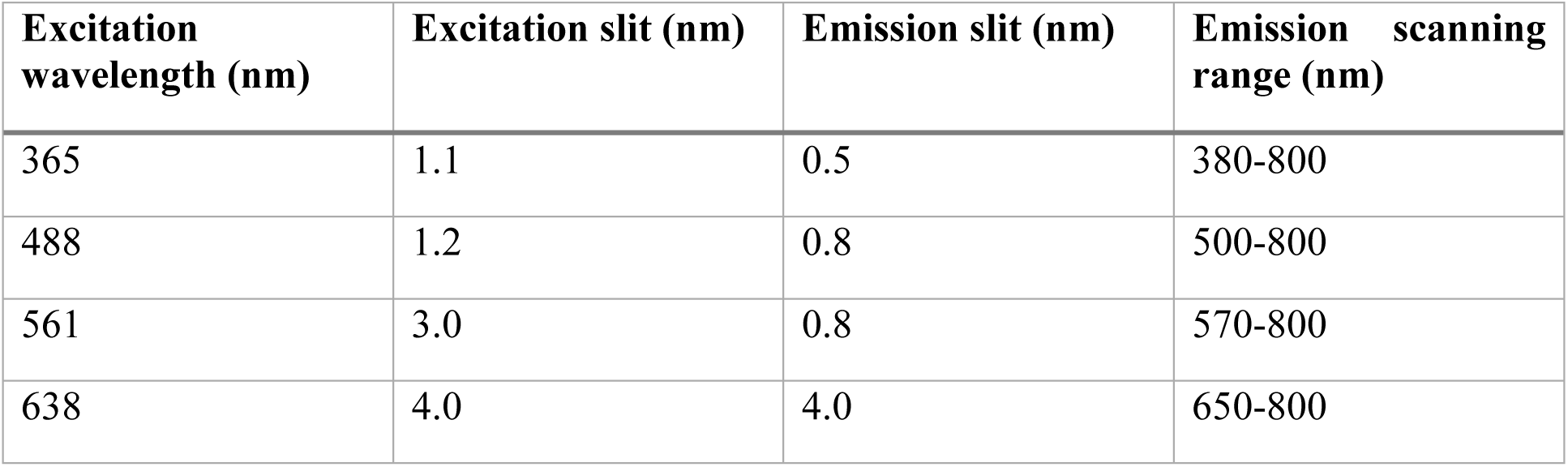
Instrumental settings used for measurement of steady-state fluorescence emission spectra of cured photoresins.

Photobleaching experiments were performed using a freshly prepared sample under continuous illumination at 488 nm with an irradiance of 54 mWcm^−2^ - measured using a calibrated power meter (PM400, Thorlabs Inc.) and sensor head (S170C, Thorlabs Inc.). The integrated fluorescent emission between 587 nm and 592 nm was measured every five minutes in 1 nm steps with a 0.5 s/step dwell time.

Raman spectra were acquired using the SRS microscope system described below. The pump wavelength was scanned across the vibrational range of interest (covering both the CH and fingerprint regions) in steps of 0.2 nm. The resulting intensity maps at each wavelength were then used to reconstruct the Raman spectra for both IP-S and IP-L.

### Dimensional calibration

Areal topography of stacked lattices and forked wedge phantoms was measured using a laser scanning confocal microscope (Lext OLS4100, Olympus), equipped with a selection of objective lenses. Topography and associated confocal intensity maps were analysed using DigitalSurf MountainsMap software (v10.3) and MATLAB (v2025b, MathWorks). The instrument scales were traceably calibrated in 20× and 50× configurations using a 50 µm VLSI step height standard and a Zygo/VLSI lateral calibration standard, and the estimated amplification coefficients used to correct dimensional outputs subject to a residual uncertainty. The Z origin was established by subtracting from the height map a least-squares plane fit to points unambiguously on the substrate surface. The topography map was also masked for low-confidence points, for example those with a low confocal intensity.

For the stacked lattice phantoms, height offset histograms were calculated from pixels within the top surface of each layer, after masking using a layer-specific minimum intensity constraint noting that level regions were locally brightest. The height of the layer was then defined as the centroid of the histogram computed for all unmasked points within 1 µm of the nominal peak value. Lateral lattice spacings were computed from the relevant layer-specific height map, as prepared and masked for height histogram analysis. For convenience the image was pre-rotated to align the lattice with XY before masked points were coerced to zero to ensure good height contrast. Cross-correlating the complete height map with a set of intersection templates – 15 µm x 15 µm patches centred on grid bar intersections - gave a 4×4 grid of local maxima location estimates. The horizontal (X) and vertical (Y) pitch was then defined as the mean X (or Y) separation between horizontal (or vertical) neighbours. These pitch measurands are in principle reproducible by end users to better than the grid uniformity. Repeating the process using the confocal intensity channel gave almost identical pitch values with similar uncertainties. A simpler alternative method based on measuring the distance between lattice bar centroids gave an equivalent X pitch, but with greater than three times the uncertainty.

To quantify the slope angle of the forked wedge phantoms the topography was acquired with the wedge of interest aligned with the X direction. After scale correction and pre-levelling via fit to good points on the substrate, a mean profile was calculated from a set of X-parallel traces corresponding to the tine centres. Least-squares lines are fitted to the points on the substrate and to the points on the slope segment, respectively, in this mean profile. The region close to the top of the slope is excluded. The slope angle was then determined from the angular difference between least squares linear fits to the substrate and wedge surface. The reported uncertainty was dominated by the reference instrument Z scale uncertainty. A linearity deviation was derived from the residuals to the fit to the slope segment. This residual profile was S-filtered according to ISO 21920-2 using an 8 µm nesting index (cut-off), and half the peak-to-valley range reported as the linearity deviation.

All dimensional uncertainties provided in the results section are given expanded to a confidence interval of approximately 95%. Reported uncertainties account for the instrument uncertainties and the precision of the phantom’s physical realization of the relevant measurand.

### Single photon excitation microscopy

Confocal fluorescence images were acquired with a Stellaris 5 laser scanning confocal microscope (Leica Microsystems) using a 20x/0.75 objective lens (HC PL APO CS2). All images were acquired with a 2 Airy unit pinhole to balance optical sectioning and photon collection efficiency. DAPI and TRITC channel images were acquired simultaneously under excitation at 405 nm and 587 nm, followed by simultaneous acquisition of FITC and Cy5 channels under excitation at 448 nm and 685 nm. Images were captured with a voxel size of 0.16 µm x 0.16 µm x 0.65 µm. Excitation irradiance and detector gain were adjusted to optimise image quality. For water immersion experiments a water droplet was added on top the phantom 20 minutes before imaging.

Light-sheet fluorescence microscopy was performed using a modified (upright) Airy beam light-sheet microscopy (M Squared Lasers) equipped with a pair of matched 10/0.3 water dipping objective lenses (UMPLFLN 10XW, Olympus), a tube lens with a focal length of 300 mm and a complementary metal oxide semiconductor camera (ORCA-Flash 4.0, Hamamatsu Photonics). To mount the phantoms within the imaging pocket formed between the nose cones of the two objectives lenses, a small section containing the phantoms was first cut from the coverslip using a diamond knife. The cut section of coverslip was then placed on top of a droplet of 1% agarose gel mounted on top of a PDMS plinth within a shallow plastic dish. Finally, the gel and coverslip were immersed in water. DAPI and FITC channel images were acquired with a single plane exposure time of 100 ms using 3% and 5% of laser power respectively. While TRITC and Cy5 channel were acquired with 200 ms exposure time using 12% and 45% of laser power respectively. All channels were acquired with voxel size at 0.387 µm x 0.387 µm x 0.5 µm. PSFs were acquired for each channel from image of multicolour fluorescent microspheres (Tetraspeck, 0.5 μm, Thermofisher) immobilized in a 1% agarose gel. Raw images were then deconvolved using the Richardson-Lucy algorithm with 100 iterations.

### Stimulated Raman scattering and two-photon excitation microscopy

SRS and TPEF microscopy images were acquired with a Leica SP8 laser scanning microscope (Leica Microsystems) coupled to a PicoEmerald-FT laser (APE) using a 25x/0.95 objective lens (HC FLUOTAR L W VISIR, Leica Microsystems) and an oil immersion condenser with an NA of 1.4 (P 1.40 OIL S1, Leica Microsystems). The PicoEmerald-FT generates two pulsed 2 ps laser beams: a 1031.2 nm Stokes beam which was spatially and temporally overlapped with a tunable pump beam. Prior to imaging, the temporal and spatial overlap of the beams were optimised using local controls and standards. The Stokes beam was modulated at 20 MHz and stimulated Raman loss signals were detected using a hybrid photodetector and lock-in amplifier (UHFLI, Zurich instruments). The power of the Stokes and pump beams was set to 36 mW and 11.7mW respectively and the pump beam was tuned to excite transitions at 1636, 1725, 2967, and 3041 cm^−1^. Background signals were removed by subtracting off-resonance images at 1679, 2695, and 3106 cm^−1^. Detector gain settings were adjusted to optimize image quality with SRS acquired using a gain of 19.2 V. TPEF signals under excitation by the 36 mW Stokes beam were detected in a second imaging channel using a photomultiplier tube with a gain of 666 V. For both SRS and TPEF modes 3D scans were acquired with a 1 μm axial step size and a pixel size of 0.91 µm.

### Scanning electron microscopy

SEM images were acquired using a ThermoFisher Apreo S2 system with a 2 kV accelerating voltage, a 0.05 nA beam current and a standard Everhard Thornley secondary electron detector. The ITO layer on the coverslip minimised charge build-up, meaning that phantoms could be imaged directly without additional coating to increase electrical conductivity.

### Cell seeded collagen scaffolds

HeLa cells expressing GFP (AKR-213, Cell BioLabs Inc) screened for mycoplasma using a universal mycoplasma detection kit (ATCC; 30-1012k) were maintained in 1× Dulbecco’s Modified Eagle’s Medium (DMEM) cell culture medium supplemented with 10% v/v fetal bovine serum (FBS) and antibiotics (gentamicin and amphotericin B) at 37 °C, 5% CO2, and 95% humidity. The cells above 80% confluency were washed (×3) with PBS and trypsinized followed by addition of serum supplemented media to eliminate secondary toxic effects of trypsin. Detached cells were spun down by centrifugation, and the excess solvent was replaced by cell growth media. Cells seeded in type I rat tail-derived collagen (5 mg/ml, Ibidi GmbH) hydrogels prepared according to manufacturer’s instructions at 1.5 mg/ml concentration. Cells were mixed in gel precursor solution at a density of 500 cells per µl and a 100 µl gel was cast in an 8-well chambered coverslip (Ibidi GmbH). Gels were placed in cell culture incubator (37 °C, 5% CO*2*, 95% humidity) for 30 min to allow for gelation before topping up with 1 × supplemented cell culture media. After 24-hours of incubation, cells were stained for F-actin (CellMask Actin Tracking Deep Red, Thermofisher) and nuclei (Hoechst 33342, Thermofisher). The actin stain was diluted 1:1000 in Opti-MEM (Gibco, Thermofisher) and the nuclear stain was diluted to 0.01 mg/ml in Opti-MEM before addition to the cell-seeded hydrogel. After 1 h at 37⁰C, 5% CO2, the sample was washed 3 times with 1x PBS and fixed for 30 min at room temperature using 4% paraformaldehyde solution (Thermofisher). The fixative was then washed off by rinsing the gel 3 times with 1x PBS.

## Author Contributions

J.J. and M.S. designed the study. B.R. and J. J. fabricated phantoms. K.M., C.W.J., D.T., J.J. and M.K. captured experimental data. P.G. prepared cell-seeded scaffolds. J.J., M.S. and C.J. performed formal analysis. M.S. wrote the first draft of the manuscript with contributions from all authors. All authors reviewed and approved the final version of the manuscript.

## ACKNOWLEDGMENTS

The authors acknowledge funding from the UK Government’s Department for Science, Innovation & Technology through the Life Sciences and Health programme of the National Measurement System.

