## Supplemental Figure 1 and Figure 2 for "Dimensionally traceable 3D microstructures for multimodal microscope calibration"

### Dimensionally traceable 3D microstructures for multimodal microscope calibration: supporting information

#### SEM micrographs showing phantoms fabricated from different photoresins at different scales

A set of forked wedge, stacked lattice and cylinder array phantom designs were fabricated at different scales; with features covering a total area from 100 µm x 100 µm to 500 µm x 500 µm. SEM micrographs (Fig. S1) indicated no obvious differences between the fidelity of structures fabricated from IP-S and IP-L photoresins. The issues with the cylinder array design noted in the main text – variable wall thickness, instability of tall cylinders and partial fabrication of short cylinders – were present all all scales. Similarly, closely-spaced internal panels tended to collapse or merge for forked wedge phantoms fabricated at small and large scales. The stacked lattice was reproduced without obvious issues across all length scales. These results suggest phantoms can be scaled to match the requirements of different microscope platforms.

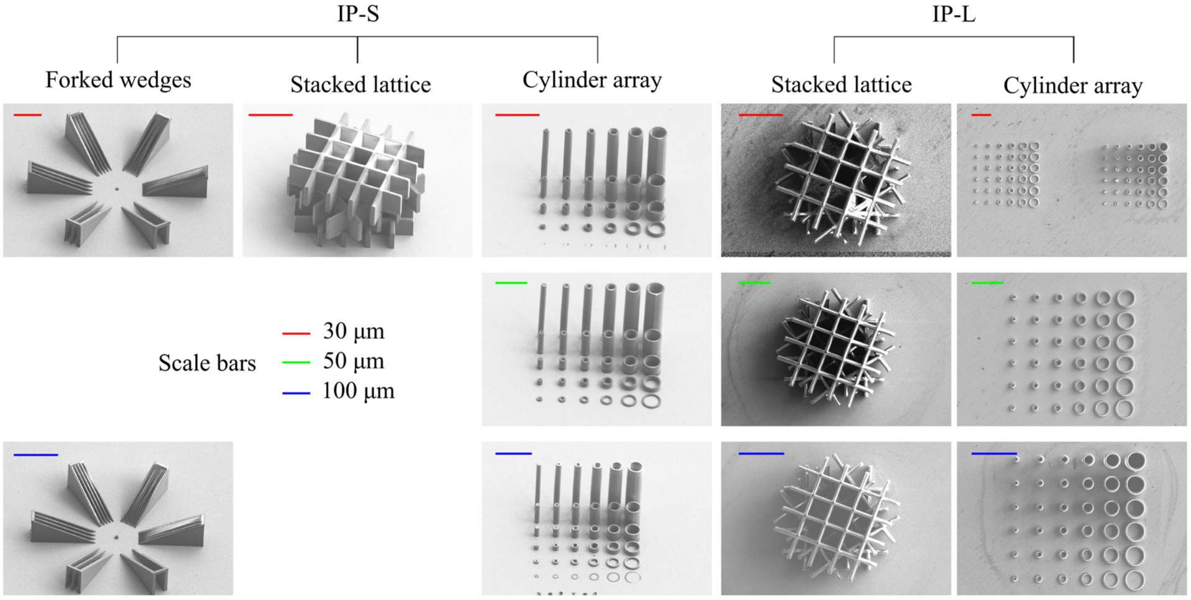


**Figure S1. SEM micrographs of IP-L and IP-S phantoms fabricated at different sizes**

#### Structural stability of phantoms

The surface topography of the same (500 µm x 500 µm) stacked lattice phatom fabricated from IP-L photoresin was measured shortly after fabrication and then again following three months of storage at room temperature. The resulting height maps (Fig. S2) indicate no signficant changes to the object structure over this period.

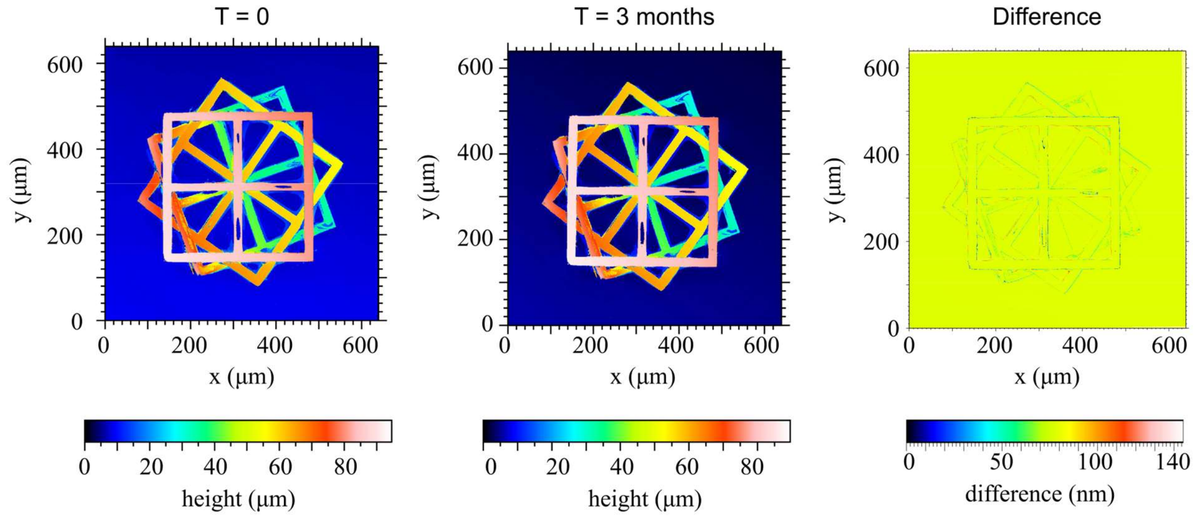


**Figure S2. Topographic calibration of a prototype stacked lattice phantom.** The same phantom was measured shortly after fabrication (left) and again after 3 months of storage (middle). The difference between calibration results (right) indicates there were no significant changes to the structure over the storage period.
